# A deep audit of the PeptideAtlas database uncovers evidence for unannotated coding genes and aberrant translation

**DOI:** 10.1101/2024.11.14.623419

**Authors:** Jose Manuel Rodriguez, Miguel Maquedano, Daniel Cerdan-Velez, Enrique Calvo, Jesús Vazquez, Michael L. Tress

## Abstract

The human genome has been the subject of intense scrutiny by experimental and manual curation projects for more than two decades. Novel coding genes have been proposed from large-scale RNASeq, ribosome profiling and proteomics experiments. Here we carry out an in-depth analysis of an entire proteomics database.

We analysed the proteins, peptides and spectra housed in the human build of the PeptideAtlas proteomics database to identify coding regions that are not yet annotated in the GENCODE reference gene set. We find support for hundreds of missing alternative protein isoforms and unannotated upstream translations, and evidence of cross-contamination from other species.

There was reliable peptide evidence for 34 novel unannotated open reading frames (ORFs) in PeptideAtlas. We find that almost half belong to coding genes that are missing from GENCODE and other reference sets. Most of the remaining ORFs were not conserved beyond human, however, and their peptide confirmation was restricted to cancer cell lines. We show that this is strong evidence for aberrant translation, raising important questions about the extent of aberrant translation and how these ORFs should be annotated in reference genomes.

## Introduction

The human reference genome has been completed with the annotation of heterochromatic regions [1] and the Y chromosome [2] by the T2T consortium. Despite this, the annotation of a final set of human coding genes is still some way from being finished. The principal reason for this is that the main reference databases [3–6] still disagree on which genes code for proteins [7, 8], although the completion of the novel CHM13 human reference, in which preliminary estimations of novel coding genes run from 2 [9] to 300 [4] has added a new layer of complexity. On top of this, there is increasing evidence from ribosome profiling experiments that a surprisingly large number of unannotated open reading frames undergo translation [10].

The initial drafts of the human reference genome [11, 12] pinned the number of protein coding genes at between 25 and 35,000. Since then, estimates of the number of coding genes have been part of a gradual downward trend [7, 13–17]. The most recent GENCODE release [3] (v46) annotates 19,411 coding genes, though there were more than 22,000 annotated in the combination of the RefSeq, UniProtKB and Ensembl/GENCODE reference sets in 2018 [7].

Recently, several high-profile large-scale ribosome profiling analyses have been published that demonstrate evidence for tens of thousands of unannotated open reading frames (ORFs) in the human genome [18, 19]. A consortium has been formed that will investigate whether these regions are likely to code for proteins [10], with one of the early papers published by members of the consortium suggests that the more than 7,000 well-supported novel ORFs they find might expand coding gene numbers by 30% [20]. Such a large increase in protein coding genes would raise the question of what constitutes a coding gene.

The paper highlights the discovery of novel ORFs that are often cited as evidence for the functional importance of short ORFs as a whole. These are *APELA*, *ASDURF*, *MIURF*, *MRLN*, *MYMX*, *POLGARF*, *TINCR*, and the as yet unnamed uORFs in *MKKS* and *SLC35A4*. What links these nine examples is not that they are short (three are longer than 100 amino acids and *POLGARF* has 260), but that they all have obvious cross-species conservation. Seven can trace their ancestry back to lobe-finned fish at least, while the other two are conserved across all mammals. However, these 9 genes are not representative of the class of novel ORFs because most novel ORFs detected in ribosome profiling experiments have little evidence of cross-species conservation.

While it is certainly true that there will be other conserved novel coding ORFs beyond the nine cited examples, we have recently detected one that overlaps the *GRIN2A* gene that is conserved across eutherian mammals and that has peptide evidence [21] for example what links the large majority of the novel ORFs detected in ribosome profiling experiments is that they usually have little or no evidence of cross species conservation. Analyses of germline variants from other coding regions with little or no evolutionary track record shows that these are generally not under purifying selection [7, 21, 22–24] and that few to none of the variants found for these regions are pathogenic [25, 26]. Human variant data strongly suggests that very few non-conserved novel ORFs will be functionally relevant, although some recently evolved functionally important coding genes certainly do exist [27].

The proteomics evidence for novel ORFs is not strong. The same two large-scale ribosome profiling analyses that detected tens of thousands of novel ORFs, failed to find much evidence for their products in standard proteomics experiments. One found peptide evidence for just seven unannotated ORFs [18], and although the other detected 541 peptides for hundreds of novel ORFs in two proteomics analyses, only five of the novel peptides (fewer than 1%) were found in both experiments [19]. The consortium themselves [10] found just 13 peptides for the novel ORFs in PeptideAtlas, with one peptide per coding gene and most of these peptides were not tryptic. The extent to which these novel ORFs produce stable proteins is not clear.

If these novel ORFs are being translated as the ribosome profiling experiments suggest, and these proteins are stable, there ought to be peptide evidence for them in proteomics analyses. So, what is happening? One explanation is technical. The smallest of the proteins and those proteins with a special amino acid composition are not amenable to detection in standard trypsin-based proteomics experiments [20], although this still leaves many thousands of undetected translated ORFs [21]. Another possible reason may be that some of the transcripts captured in ribosome profiling experiments are not translated, for example due to control mechanisms at the level of the ribosome [28]. If translated, it is also possible that many of these peptides are rapidly degraded [29]. There is certainly evidence for some degradation in the ribosome profiling-based analyses [18,19], since there are plenty of novel ORF peptides in proteasome-derived human leukocyte antigen class 1 (HLA-I) proteomics experiments [20, 30]. The final possibility is that the novel ORFs are translated rarely or in low quantities. If this was happening, a single proteomics analysis would be unlikely to find much evidence, but multiple large-scale proteomics experiments might be able to provide some support.

Here we carry out a manual analysis of the novel ORFs detected in the PeptideAtlas database [31]. PeptideAtlas is a database that maps annotated and predicted proteins to thousands of proteomics experiments. PeptideAtlas interrogates spectra from many varied proteomics experiments and its search database contains many predicted proteins that are not produced by genes in the reference databases. The combination of these three features (and manual curation possibilities) means that PeptideAtlas is a potential source of evidence for protein coding genes still missing from the reference annotations.

Instead of starting with a list of transcripts from ribosome profiling experiments, we downloaded all PeptideAtlas peptides with the aim of discovering protein coding regions that the Ensembl/GENCODE reference might have missed. We found evidence for hundreds of regions that are not annotated as coding in the GENCODE version of the human reference gene set. Although most of the non-reference peptides mapped to alternative variants of known proteins, there was convincing evidence for proteins from 34 novel coding ORFs, most of which have little cross species conservation. We found peptide evidence for seven of the short ORFs predicted by ribosome profiling experiments [18,19], and we believe that five of these represent coding ORFs that are novel to Ensembl/GENCODE. We predict that other two proteins are the result of aberrant translations.

## Methods

### Generating a list of novel peptides from PeptideAtlas

We downloaded peptides from the January 2023 build of the human PeptideAtlas repository. Peptides are pre-mapped by the PeptideAtlas Trans-Proteomic Pipeline (TPP, 32), a suite of locally installed tools. The TPP maps spectra from large and small-scale proteomics experiments to the PeptideAtlas human protein search database. We analysed the peptides from PeptideAtlas because the search database includes proteins from a wide range of sources and because TPP has stringent statistical validation at the peptide and protein level.

The search database (THISP, 33) is made up of sequences from the UniProtKB [6], NextProt [34], and RefSeq [5] databases, as well as likely contaminants, microbes and many non-reference peptides provided by contributors. The version of THISP used in the January 2023 build had 341,040 sequence distinct entries, Sequences from contributors provided the largest number of entries to the database. Contributor sequences are mostly protein sequences culled from large-scale experiments that might be protein coding. More than two thirds (67.8%) of the novel ORFs identified by Chen *et al* [18] and van Heesch *et al* [19] were annotated in the January 2023 build of the THISP database.

The TPP validated 3,489,945 peptides from the January 2023 build and mapped these peptides to 62,245 protein entries. Peptides identified by the pipeline were mapped to a single entry in the PeptideAtlas file. For our analysis this had two drawbacks. Firstly, peptides might map to more than one protein entry, and many do because many of the entries in the THISP database have common protein sequences. Secondly, peptides that might have been produced from a single transcript are often mapped to two or more distinct entries (due again to the abundant regions of common protein sequences in the THISP database).

To detect which peptides mapped to proteins outside of the Ensembl/GENCODE reference set [3, 4], we remapped the 3,489,945 peptides validated by the Trans-Proteomic Pipeline to the GENCODE v43 reference set of proteins [Figure 1]. The GENCODE v43 reference set was downloaded from the GENCODE website. We excluded all peptides that were not fully tryptic. Peptides that mapped to GENCODE proteins were excluded from the remainder of the analysis. We also excluded decoy peptides, and peptides that mapped to common contaminants and to microbes.

**Figure 1.**
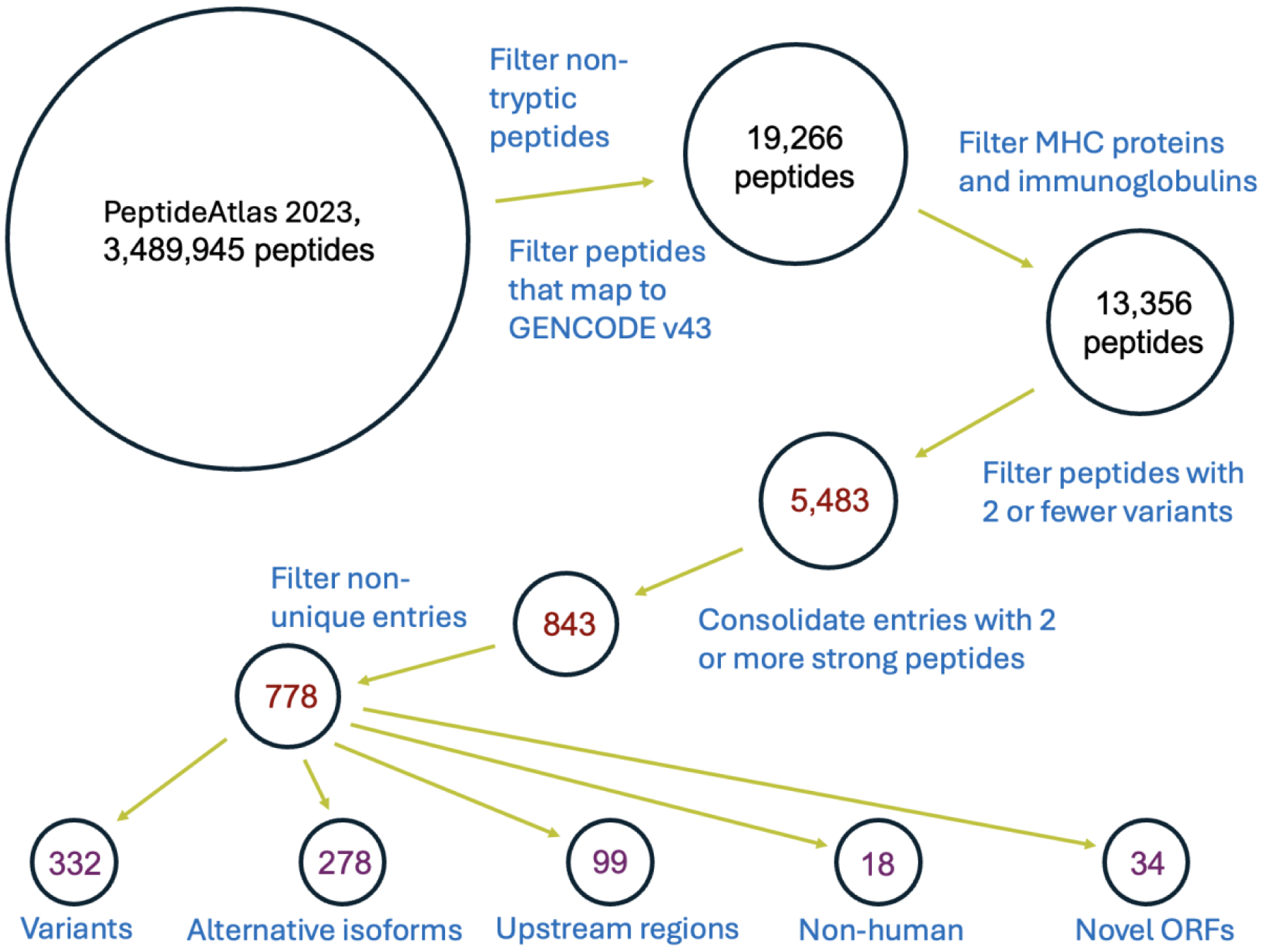
Workflow. The analysis of PeptideAtlas peptides that did not map to proteins in the Ensembl/GENCODE reference set and the five largest classes of “missing” proteins that were identified.

After filtering, we were left with a total of 19,266 tryptic peptides that did not map to GENCODE genes. A further filtering step removed 5,910 novel peptides that we knew mapped to UniProtKB immunoglobulins or MHC proteins. We know that these proteins will differ in amino acid sequence from the GENCODE entries from the reference set. This left us with 13,356 novel peptides.

### Defining strong peptides

We know that there are many probable pseudogenes in the THISP search database [7], and we know that many of the pseudogenes have peptides that differ from their parent genes by just one or two amino acids. Proteomics experiments do not always correctly distinguish canonical peptides from theoretical peptides pseudogenes, largely because of single amino acid variations (SAAVs) and post-translational modifications [33]. Proteins often have many natural commonly occurring SAAVs. One way to get around these problems would be to just remove any peptide that was a SAAV of a GENCODE protein. However, this would make it impossible to uncover *bona fide* novel coding genes that are close homologues of other coding genes.

The downloaded list of PeptideAtlas peptides includes the number of times the peptide has been observed across the experiments interrogated in PeptideAtlas. The more observations, the more common the peptide. We made use of the number of observations for each peptide to score peptides that were just one or two amino acids different from peptides that mapped to GENCODE proteins. For each peptide we summed the observations for all peptides that differed by two or fewer amino acids. These we termed double amino acid variant (DAAV) peptides. The observation score for each peptide was then the number of observations for the peptide divided by the sum of observations from all DAAV peptides. We regarded peptides with observation scores above 0.9 as strong peptides. Peptides that had no DAAV peptides automatically had observation scores of 1. These strong peptides are referred to in the paper as strong discriminating peptides (SDPs).

### Manual curation of the entries supported by strong discriminating peptides

We found 5,483 SDPs in PeptideAtlas that did not map to GENCODE reference set proteins. These peptides mapped to 3,774 distinct non-GENCODE entries. Most entries were supported by a single SDP, but 843 entries had two or more mapping SDPs [Figure 1].

We manually curated the novel peptides for the 843 entries that had two or more SDPs that did not map to the GENCODE annotation. Manual curation attempted to ascertain where the peptide mapped, whether the peptide was most likely to be a variant of a known protein sequence (despite our filtering for amino acid variants some variants still got through the filters), whether it was evidence for a novel splice isoform or a novel ORF in an untranslated region of a gene, or whether the peptide supported a whole new ORF that was not annotated in GENCODE.

Since peptides from the same gene often map to more than one entry in the THISP database, we combined entries where the peptide data supported the same protein product. Peptides for the LINE-1 ORF1 protein mapped to nine distinct THISP entries, for example, so we combined the evidence into a single entry during the manual curation process. After combination the number of PeptideAtlas entries with at least two strong novel peptides was 778.

## Results

### Most novel peptides are most likely to be variants, modifications or false positives

Most of the 13,356 novel peptides identified by PeptideAtlas were single or double amino acid variants of peptides from GENCODE proteins, so were likely to be either amino acid variants or post-translational modifications of known peptides. In this analysis we used the PeptideAtlas observation count to score all peptides and used only strong discriminating peptides (SDPs), those that were either substantially different from GENCODE peptides or that were similar GENCODE peptides but detected in much higher quantities (see methods section), to search for novel genes and coding regions.

### Many apparently novel entries in PeptideAtlas are not novel

In total, we defined 5,483 of the 13,356 peptides as strong. We carried out an in-depth curation of those PeptideAtlas entries that had at least two SDPs, 778 entries in total (see methods section). We considered each of these novel protein coding ORFs on its merits. We found that many of the novel entries that had the support of SDPs were still most easily explained as variants or post-translational modifications of known proteins. This is largely because amino acid variants involving lysines or arginines can produce very different peptides due to trypsin cleavage. There were also peptides with insertions or deletions of one or more amino acids in low complexity regions that were most easily explained as variants.

Some entries were particularly complicated to decipher. For example, the novel protein from UniProtKB-Trembl, A0A0F7G8J1, is supported by 24 uniquely mapping PeptideAtlas peptides, which strongly suggests that it is produced from a novel ORF. This entry is a homologue of plasminogen (*PLG*) [Figure 2], to which it has 80.4% identity. There is considerable peptide evidence for the canonical plasminogen protein in PeptideAtlas, so it is conceivable (but unlikely) that all 24 peptides that map to A0A0F7G8J1 could be variants of plasminogen peptides.

**Figure 2.**
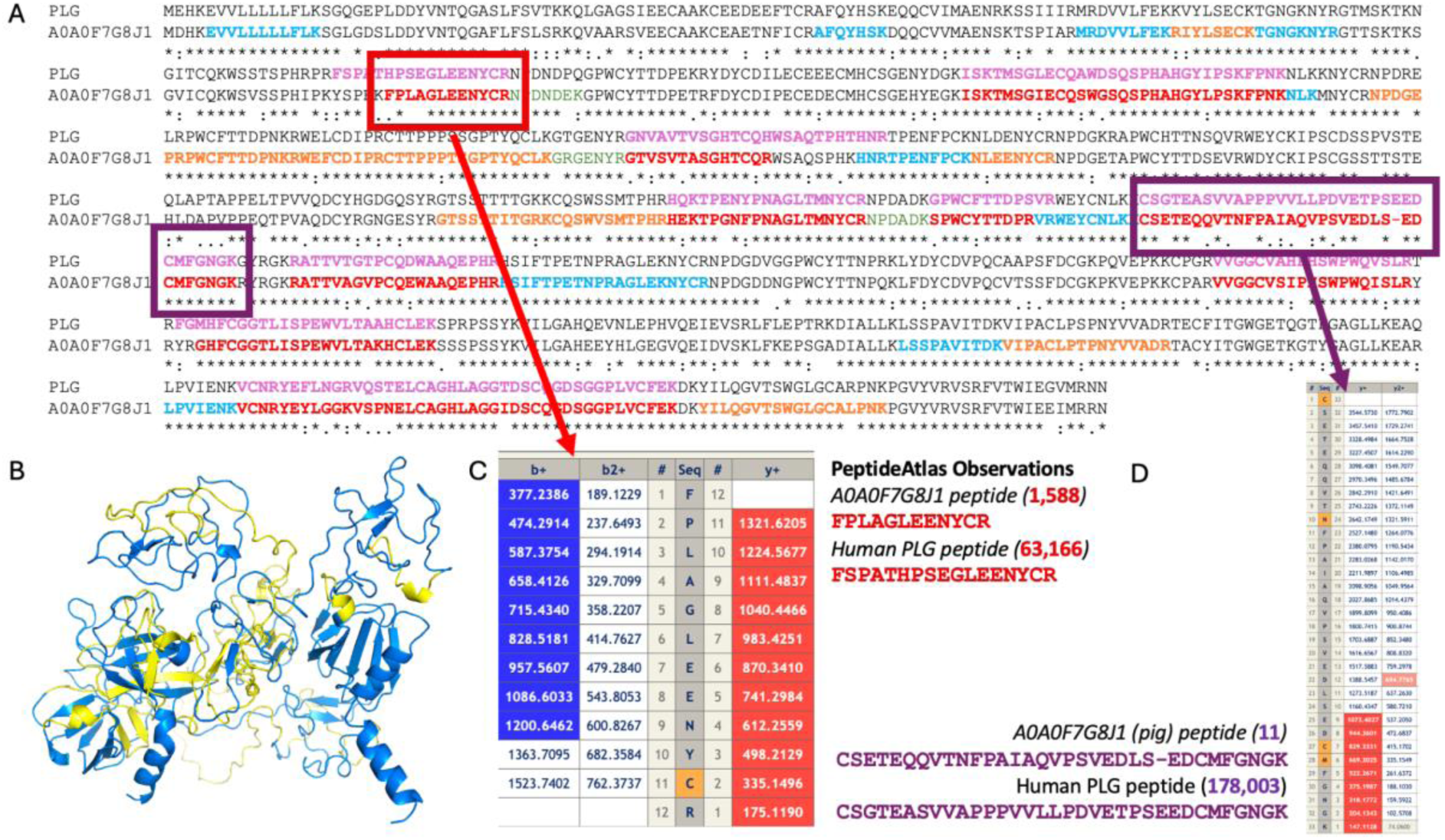
Analysing the SDPs mapping to a plasminogen homologue, A0A0F7G8J1. A. The alignment between A0A0F7G8J1 and human plasminogen with the PeptideAtlas peptides in bold and coloured by type. Red peptides are SDPs, orange peptides distinguishing peptides, but not SDPs and blue peptides are non-discriminating peptides. The tryptic peptides equivalent to the SDPs in plasminogen are marked in pink. B. The Alphafold [34] model of A0A0F7G8J1 with detected peptides marked in yellow. C. Part of the PeptideAtlas peptide spectrum match for the A0A0F7G8J1 peptide FPLAGLEENYCR showing that the y-series (in red) is a perfect match. D. Part of the peptide spectrum match for the A0A0F7G8J1 peptide CSETEQQVTNFPAIAQVPSVEDLSEDCMFGNGK showing that the y-series (in red) only matches the N-terminal residues that are identical in the equivalent *PLG* peptide.

Twelve of the 24 peptides are only one or two amino acids different from the plasminogen peptides that PeptideAtlas detected and have many fewer observations. But the other 12 discriminating peptides were sufficiently dissimilar from the equivalent plasminogen peptides to be classed as SDPs. If the 12 peptide-spectrum matches (PSMs) were good quality, this would be more than enough to support the presence of protein A0A0F7G8J1 because it is hard to imagine how amino acids variants and PTMs could explain so many large differences.

However, the PSM evidence is mixed. For example, the A0A0F7G8J1 peptide “FPLAGLEENYCR” has 1,588 observations, is very different from the equivalent peptide in plasminogen, and has very good looking PSM evidence [Figure 2]. This peptide supports a novel ORF for this protein. On the other hand, other peptides do not have such good PSM. The peptide “CSETEQQVTNFPAIAQVPSVEDLSEDCMFGNGK” has just 11 observations (compared to 178,003 for the equivalent plasminogen peptide) and PSMs that only supports the final 9 amino acid residues. These final 9 amino acid residues are identical between the A0A0F7G8J1 peptide and the equivalent plasminogen peptide, so these PSM do not supports the theory that the A0A0F7G8J1 protein is translated.

In fact, A0A0F7G8J1 is 99.4% identical to pig plasminogen (UniProtKB entry P06867) and does not map anywhere in the human genome. It is possible that pig plasminogen or something similar may be a contaminant that has not yet been included in the PeptideAtlas database.

A total of 322 entries were classified as most easily explained as amino acid variants or PTMs of canonical human proteins. Many of these entries are actin-like proteins. There are many actin-like entries in the PeptideAtlas THISP search database [33], partly because there are multiple actin pseudogenes in the human genome. Canonical actin proteins are highly expressed and highly conserved; seven of the ten entries with most PSM in PeptideAtlas are actins. Peptides that map to actin-like entries from the THISP search database have many fewer supporting PSM, and it is highly likely that the spectra that map to these actin-like entries belong to canonical actin peptides and are mis-mapped to the actin-like pseudogenes instead. The mis-mapping may be because of modifications or variants, or simply because of false positive match from a low-quality spectrum. Like actin, other genes and gene families, including haemoglobins, ribosome proteins and *GAPDH*, also have multiple erroneously identified pseudogenes.

### Multiple non-human proteins detected in human proteomics experiments

HEL-109, a novel epididymis luminal protein, is identified with 13 uniquely mapping tryptic peptides in our analysis and in the January 2024 version of PeptideAtlas there are spectra for 24 distinct fully tryptic peptides. This novel 711 amino acid protein [Figure 3] is annotated by UniProtKB and appears to be a homologue of tropomyosin; the C-terminal region has more than 60% identity to one of the human *TPM3* isoforms.

**Figure 3.**
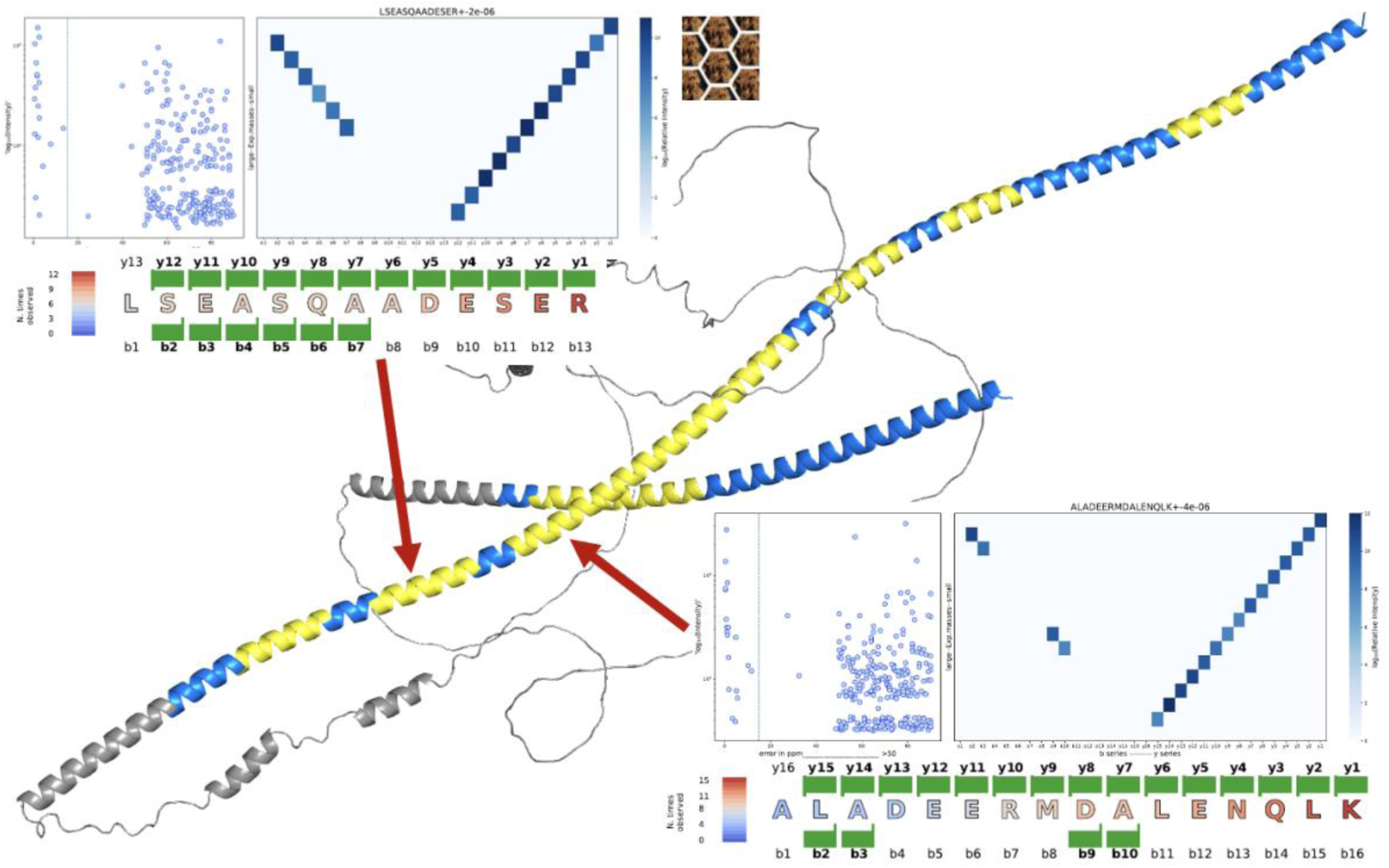
Validating peptides for *Drosophila melanogaster* “HEL-109”. The AlphaFold model of UniProtKB protein HEL-109 (V9HVX8) with the detected peptides mapped in yellow. HEL-109 is really the product of an extended *Drosophila melanogaster* alternative transcript (Tm1-RE). The regions of V9HVX8 that are identical to the *Tm1* principal isoform [36] are coloured blue (where no peptide matches) and yellow (where peptides from PeptideAtlas map). The structure is coloured grey for those regions of V9HVX8 that are substantially different from the *Tm1* principal isoform. There are no peptides for these 426 residues. Two of the peptides (positions shown by red arrows) were analysed using VSeq [37], and the VSeq output shows that these two peptides could not be mistaken for other peptides. Similar results were found for almost all the detected HEL-109 peptides.

However, HEL-109 is not a human protein, it is a minor isoform of *Drosophila melanogaster* tropomyosin, *Tm1*. The protein comes from a CDS deposited in GenBank [40] in 2008 (EU668323) that has been incorporated into UniProtKB TrEMBL. TrEMBL proteins form part of the THISP search database. The Flybase [38] Tm1-RE transcript almost certainly does not produce a functional 711 amino acid protein, it has TRIFID [39] score of 0.263 against 1.0 for the principal transcript Tm1-RG. All 24 SDPs map to the protein produced by Tm1-RG.

How did a fruit fly protein come to be identified by so many uniquely mapping peptides in human mass spectrometry experiments? One possibility is that the search engine simply mistook the *Drosophila melanogaster* peptides for human peptides. However, we have analysed the spectra for the 24 peptides and 21 of them are supported by at least one peptide-spectrum match that has a perfect y-series [Figure 3]. The identifications of the *Drosophila melanogaster* peptides are not erroneous.

Another possibility is that human tropomyosin genes have single amino acid variants or post-translational modifications that mimic their *Drosophila melanogaster* counterparts. However, almost all the 24 *Drosophila melanogaster* peptides have three or more amino differences from the human counterparts. Human tropomyosin genes have few non-synonymous variants and those missense variants that are known do not correspond to the amino acids that differ between the human and fruit fly peptides. These are not variants.

Post-translational modifications in human tropomyosin are less well annotated, but even here it is hard to imagine that they could be the reason why fruit fly proteins were identified. With 21 of the 24 peptides supported by PSMs with perfect y-series, there would need the perfect combination of multiple variants and post-translational modifications in the human peptide to confuse the PeptideAtlas search engine. And this needs to have happened not just for one peptide, but for all 21 peptides. This explanation is clearly impossible.

So, only one possibility remains. The PeptideAtlas search has identified a *Drosophila melanogaster* protein in human experiments. Ignoring explanations that belong in science fiction, it means that experiments were contaminated in some way with fruit fly proteins.

Twenty three of the 24 peptides were detected in only four proteomics experiments. The final peptide detected had only a single PSM and may be a false positive identification. The four experiments covered a range of unrelated tissues, including ovarian tumours, placenta, and peritoneum. Each experiment identified between 7 and 10 distinct fruit fly peptides and all 21 perfect y-series came from these four experiments. These experiments were all somehow contaminated with insect proteins.

PeptideAtlas also identifies multiple peptides for five mouse proteins that are misannotated as human in UniProtKB Trembl. We analysed these five proteins in detail and concluded that the identification of these mouse peptides was almost certainly due to cross-contamination of mouse proteins in human experiments. A small number of proteomics experiments identified multiple peptides for three or four of the different mouse proteins, but as many as 50 or 60 distinct human proteomics experiments had evidence for at least one of these mouse proteins.

There are also proteins from invasive bacteria and virus that are annotated as human by UniProtKB, though it is easier to imagine how these came to contaminate human proteomics experiments. For example, the novel protein with the seventh highest number of SDPs was derp12 (Dermal papilla derived protein 12), which a BLAST [41] search shows is a dihydrolipoyl dehydrogenase from *Mycoplasma hyorhinis*, a swine pathogen sometimes found on human skin [Figure 4]. Entries tagged as human by TrEMBL and validated by the PeptideAtlas human build include proteins from E. Coli, pseudomonas and papillomavirus, that clearly should not be annotated as human proteins. These proteins could be added as part of the “microbe” section of the PeptideAtlas THISP database.

**Figure 4.**
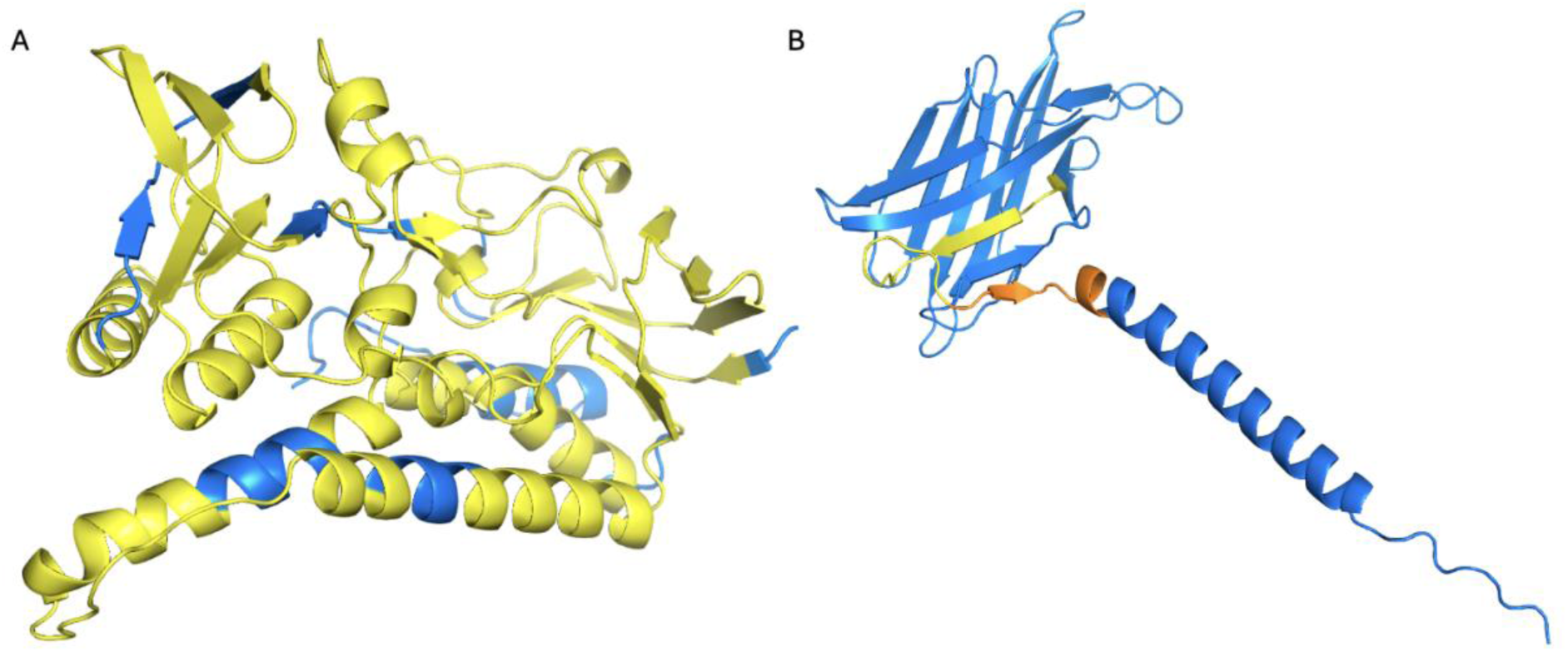
Identified peptides for Derp12 and the *LAMP3* alternative isoform. A. The AlphaFold predicted structure for Derp12 painted yellow to indicate the peptides detected. B. The AlphaFold predicted structure for the *LAMP3* alternative isoform (XP_005247417.1) showing the alternative C-terminal helix. The region with PeptideAtlas peptides is painted yellow where it overlaps the *LAMP3* region common between the alternative and principal isoforms and orange where it overlaps the alternative C-terminal helix.

In total, we find at least two PeptideAtlas peptides for 18 non-human proteins that are annotated as human in UniProtKB. This result should be somewhat concerning for the UniProtKB TrEMBL database, since the misclassified proteins that PeptideAtlas detects are likely to only be the tip of the iceberg.

### Non-reference proteins

Four of the ten novel entries with most supporting evidence were non-reference proteins. These are proteins that are not included in the Ensembl/GENCODE reference set. Three of the entries in the top ten of SDP support correspond to the loss of function genes, *GBA3*, *GPATCH4* and *PNLIPRP2*. These genes have high frequency alleles with frameshifts or stop codons. However, the proteins are clearly expressed in a large part of the population at least. *GPATCH4* has a high frequency frameshift allele that produces a different, extended C-terminal. PeptideAtlas has considerable peptide evidence for the C-terminal produced from the high frequency frameshift and none from the original C-terminal. GENCODE does not currently include full length loss of function proteins in its downloadable proteome.

### LINE-1 ORF1

The entry with most SDP support (65 peptides) is the LINE-1 retrotransposable element ORF1 protein. The LINE-1 element that produces both this protein and an ORF2 protein that was practically undetected in PeptideAtlas [42] is present in multiple copies in the genome. The UniProtKB LINE-1 ORF1 protein maps to 37 different sites in the genome and there are more than a thousand full length LINE-1 ORF1 open reading frames that could produce ORF1 proteins with 95% or more identity to the UniProtKB protein. LINE-1 ORF1s make up more than a million bases and the sheer scale means that coordinate-based reference sets like Ensembl/GENCODE and RefSeq cannot annotate LINE-1 ORF1 as coding.

As many as 100 LINE-1 elements are still active [43]. These are responsible for retrotransposition of processed pseudogenes and SINE Alu and SINE Ventr Alu elements and are suppressed in normal tissues but become activated in tumours [44]. It is noticeable that the vast majority of observations for the 65 peptides in PeptideAtlas come from cancer cell lines. Given their clinical importance and abundance in cancer tissues [45, 46], it seems that diagnostic proteomics experiments ought to include the LINE-1 ORF1 protein in their search databases, something that could not happen with the RefSeq and/or Ensembl/GENCODE gene sets alone.

### Unannotated alternative splice isoforms

PeptideAtlas has proteomics support for 278 alternative splicing events that are not annotated in GENCODE. One example is a novel C-terminal in *LAMP3* [Figure 4B] produced from an alternatively spliced tandem duplicated exon that can be traced back to vertebrates [47]. However, none of the 278 predicted splice isoforms have yet been validated by GENCODE manual curators, and we did not analyse any of the PSM for this entry class, so we cannot rule out the possibility that some of these entries may be variants or false positive identifications, or that some may be the result of aberrant or noisy translation.

### Translation of upstream untranslated regions

There have been a number of recent publications highlighting translation from the untranslated regions (UTR) of protein coding genes, in particular translation from the 5’ UTR [21, 48, 49]. PeptideAtlas has multiple SDPs for 99 novel entries derived from the translation from 5’ UTR regions. Most of these translated upstream regions are in the same reading frame as the main transcript and would result in N-terminal extensions of the main protein isoform. We also found evidence for the translation of uORFs (ORFs that begin and end upstream of the canonical ATG), and uoORFs (ORFs that begin upstream of the canonical ATG and read through to coding exons, but in a different frame). Since these ORFs are found in the 5’ UTR regions of coding genes, we did not consider these regions as novel coding genes.

Most of these translated upstream regions have little cross species support and may be products of an aberrant or noisy translation initiation process [21, 49]. As with the alternatively spliced isoforms, these UTR decorations have not yet been validated by GENCODE manual curators, and we did not analyse the PSM for these regions, so some may be false positive identifications.

### Genes that are not annotated as coding in the GENCODE reference set

As well as non-reference genes, alternative isoforms and translated upstream regions, PeptideAtlas also has SDP support for 34 possible coding genes that would be novel to the Ensembl/GENCODE gene set. We labelled these genes as coding, aberrant or unplaced after manual curation of the data associated with each novel coding region.

The 16 likely coding genes are listed in Table 1. Two of these we had detected in an earlier analysis [9]. *WASHC1* (12 SDPs) and *GPRIN2L* (3 SDPs) are both part of the new CHM13 Assembly. Neither the correct *WASHC1* gene nor *GPRIN2L* were annotated in UniProtKB, nor in the Ensembl/GENCODE or RefSeq reference assemblies that are based on the GRCh38 assembly.

**Table 1.** Predicted likely coding genes. The 16 genes supported by PeptideAtlas that are not in the Ensembl/GENCODE reference set and are likely to be protein coding.

| Gene | Strong Peps | UniProt | Annotated | Decision |
| --- | --- | --- | --- | --- |
| *ANKRD26P1* | 9 | Proteome | Pseudogene | Coding |
| *C5orf60* | 16 | Proteome | Pseudogene | Coding |
| *EP400P1* | 6 | Proteome | Pseudogene | Coding |
| *ERVFRD-2* | 3 | TrEMBL | Retrovirus | Coding |
| ETD-L (Q3ZM62) | 3 | TrEMBL | Pseudogene | Coding |
| *FAM183BP* | 6 | Proteome | Pseudogene | Coding |
| *GPRIN2L* | 3 | - | NA | Coding |
| *LIPT2-AS1* | 3 | TrEMBL | Pseudogene | Coding |
| *LNCPRESS1* | 2 | - | Retrovirus | Coding |
| *LOC107986768* | 4 | - | Retrovirus | Coding |
| *MSL3P1* | 9 | Proteome | Pseudogene | Coding |
| *PLAC4* | 3 | TrEMBL | Retrovirus | Coding |
| *TSPY26P* | 2 | Proteome | Pseudogene | Coding |
| *TXLNGY* | 5 | Proteome | Pseudogene | Coding |
| *WASHC1* | 10 | - | NA | Coding |
| *ZNF840P* | 12 | Proteome | Pseudogene | Coding |

### Novel ORFs from predicted pseudogenes

There are multiple peptides in PeptideAtlas for ten genes annotated in UniProtKB as part of the human reference proteome, but that were annotated as pseudogenes in both Ensembl/GENCODE and RefSeq. They are *ANKRD26P1* (9 SDPs, Figure 5A), *CFAP144P1* (6 SDPs), *EP400P1* (6 SDPs), *MSL3P1* (9 SDPs), *MYH16* (11 SDPs), *PMS2CL* (3 SDPs), *POM121L1P* (3 SDPs), *TSPY26P* (2 SDPs) *TXLNGY* (5 SDPs), and *ZNF840P* (12 SDPs). *ANKRD26P1* [Figure 5A], *EP400P1*, *MSL3P1* [Figure 5B] and *MYH16 are* highly truncated versions of their parent genes. We believe that 7 of these genes are likely to be coding genes.

**Figure 5.**
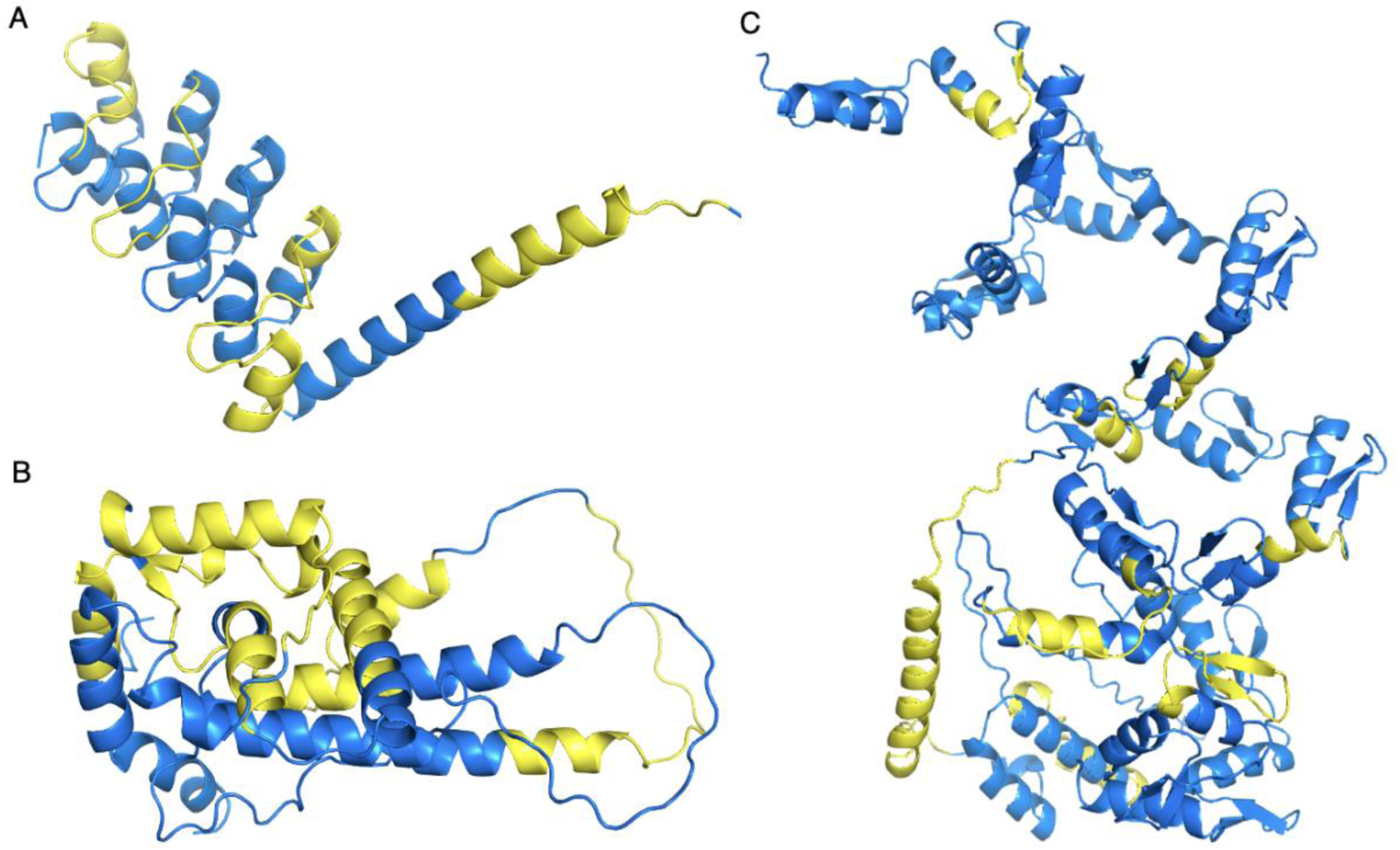
Discriminating peptides detected for *ANKRD26P1*, *MSL3P1* and *ZNF840P*. All structures predicted by AlphaFold for the UniProtKB database. Detected peptides mapped in yellow. A. *ANKRD26P1* protein with the disordered N- and C-termini removed. B. *MSL3P1* protein from the downstream ATG with the large, disordered loop removed for clarity. C. *ZNF840P* protein.

There were nine SDPs for *MSL3P1*, all of which are substantially different from the equivalent in the parent gene, *MSL3*. This is a primate duplication which has lost the N-terminal thanks to a frameshift [Figure 5B]. The peptide evidence suggests that *MSL3P1* uses a downstream ATG that would bypass the tudor knot domain in the equivalent region of *MSL3*. In addition, great apes have a different C-terminal because of a frameshift that affects the last two codons. Despite the recent changes, alignments suggest that *MSL3P1* is under protein-coding selection across all primates.

Although RefSeq has recently made *MSL3P1* coding, there is some evidence to suggest that this gene product may be the result of aberrant translation. Almost all the peptide evidence is observed in cancer cell experiments, with just a small number of unconvincing spectra found in lung. All four recent papers published on *MSL3P1* suggest that it may be an oncogene [50].

Peptides for *EP400P1* are mostly detected in cancer cell experiments, but there are also many observations in normal tissues, particularly testis. As with *MSL3P1* there are also four recent publications for *EP400P1*, and these suggest a biological role for this misannotated pseudogene [51]. *TSPY26P* has few supporting peptides but does have good conservation evidence. It is a conserved single exon gene that is under coding selection across primates, and that has orthologues in mammalian species including squirrels, whales and dolphins, and bats. Peptides were found for the N-terminal end of the protein that is almost entirely different from any paralogue.

*ZNF840P* has the most convincing evidence, though it may have a different N-terminal from the one annotated in the UniProtKB protein [Figure 5C]. *ZNF840P* would be a simian duplication that has evolved considerably. It has little more than 40% identity to other zinc finger proteins and is perhaps most similar to *ZNF780B*. The start and stop codons and the frame are conserved among old and new world monkeys. All PeptideAtlas peptides for *ZNF840P* were detected in oocytes, which fits with the RNAseq data.

The evidence *for MYH16* is conflicting. Superficially, 11 SDPs in both cancer cell experiments and in heart and muscle tissues is strong peptide support. However, the heart and muscle PSMs are poor and do not support the *MYH16* peptides. Only the PSMs from the cancer cell line experiments unequivocally support *MYH16*. In isolation, the peptide data suggests that this is an aberrant coding gene. *MYH16* is coding in primates and is specifically expressed in jaw muscles, but the human protein would have to start from a downstream start codon because of a stop codon in exon 18. In addition, human *MYH16* has lost the 5’ splice signal of exon 26. Repurposed pseudogenes do exist, but *MYH16* has no support beyond peptides in cancer cell lines. It is upregulated in lung adenocarcinoma and other cancers [52].

Finally, it is hard to imagine how *PMS2CL* could be a *bona fide* protein coding gene. It has three SDPs and all are at least three amino acids different from the parent gene. However, *PMS2CL*, currently a pseudogene, would produce a protein that would be truncated at both ends. It is also only conserved in human. The PSM found for *PMS2CL* are only found in cancer cell lines, so this may be a protein that is expressed under aberrant conditions [Table 2].

**Table 2.**
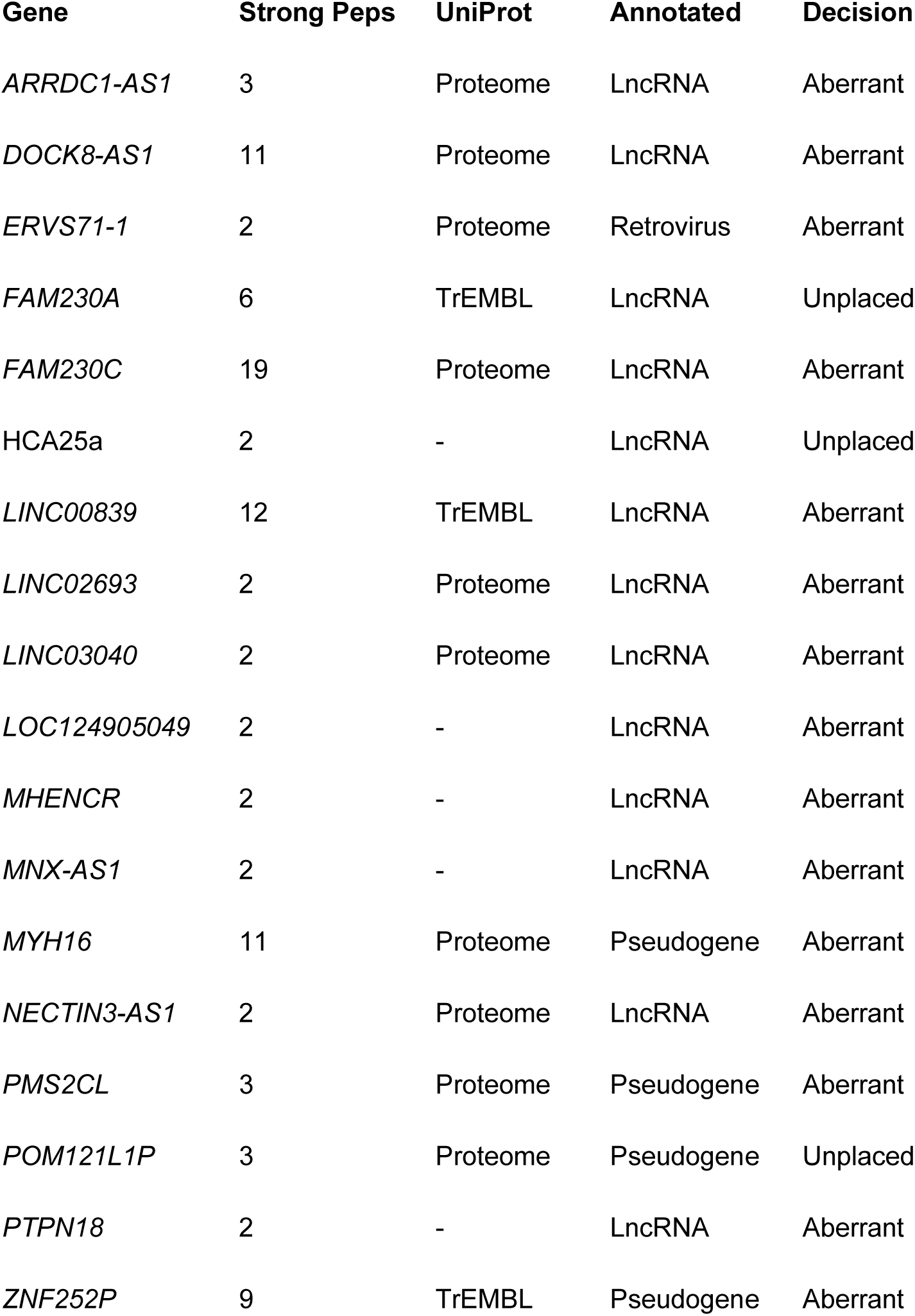
Predicted aberrant and unplaced coding genes. The 18 genes with peptide support in PeptideAtlas that are not in the Ensembl/GENCODE reference set and that are either likely to be aberrant coding genes or do not map to the genome

### Proteins from long non-coding genes

We found peptides for seven genes that were part of the UniProtKB reference proteome, but that were classified as non-coding in both Ensembl/GENCODE and RefSeq reference sets. *C5orf60* was supported by 16 SDPs, *DOCK8-AS1* by 8, *ARRDC1-AS1* by 3, and *ERVS71-1*, *LINC02693*, *NECTIN3-AS1*, and *LINC03040* by 2 SDPs each.

An eighth entry was included in the UniProtKB proteome but is not part of the GENCODE reference set. It was supported by 19 SDPs in our analysis. The protein sequence of Q9UF83 is made up almost entirely of nine amino acid repeats and there are many highly similar repeat regions of the genome, especially on chromosomes 13 and 22. Despite the overwhelming peptide evidence in PeptideAtlas, it is noticeable that all 19 peptides were detected in cancer cell lines.

Of the eight ORFs, we believe that only *C5orf60* is coding. Even though *C5orf60* is annotated as “TEC” in Ensembl/GENCODE and non-coding RNA in RefSeq, it has coding ancestry because it is a truncated copy of parent gene *SPATA31E1*. All 16 SDPs were detected in testis.

If *DOCK8-AS1* were a coding gene, it would be a *de novo* emergence from a non-coding sequence. Multiple stop codons and frame shifts means this ORF would not be translated in any other species. The peptides are all supported by their spectra, and while most are found in kidney cancer cell lines, one is found in kidney tissue. The kidney-specific expression fits with the RNAseq data from GTex [53], which shows that kidney is the most common tissue for *DOCK8-AS1*. There is no doubt that this protein is translated, but the translation is likely to be aberrant. Not only would *DOCK8-AS1* have to be a novel human-specific coding gene, but it also overlaps the first coding exon of *DOCK8* on the other strand. The expression of *DOCK8-AS1* has been shown to be related to survivability in renal cancers [54–55].

*LINC03040* was protein-coding for many years, but the CDS was removed because it had little or no evolutionary support. However, there are multiple distinguishing peptides for this gene, including ten non-tryptic peptides that we filtered out at the start of the automatic analysis. These ten peptides and the only strong tryptic peptide with a good-looking PSM were all detected in HLA-I experiments. So, *LINC03040* does have considerable evidence for translation, but the most reliable peptides are detected exclusively in HLA proteomics experiments, most of which were carried out on cell lines. We still do not have a clear idea why some peptides are detected in HLA experiments, but not in normal tissue, but *LINC03040* seems most likely to be an aberrant rather than a functional coding gene. It was recently suggested that *LINC03040* is an oncogene in colon carcinoma [56].

### Possible novel genes annotated only in the TrEMBL database

Eight ORFs with peptide evidence were novel to GENCODE but found in the catch-all UniProtKB TrEMBL database. *LINC00839* was supported by 12 SDPs, *ZNF252P* by 9, *FAM230A* by 6 and Q3ZM62 by 5. Three other cases (*ERVFRD-2*, *LIPT2-AS1*, and *PLAC4*) were supported by three SDPs each, and HCA25a by just two peptides.

The lncRNA *LIPT2-AS1* has 42% identity to the *JRK* gene, which is a Tigger transposon domestication event, but the *LIPT2-AS1* protein would not have the DDE endonuclease domain central to the *JRK* protein. The Cactus 470 mammal alignments [57] show clear protein coding conservation right across all primates for *LIPT2-AS1* and UniProtKB annotates highly similar proteins for rabbit, naked mole rat, beaver, green anole and bamboo shark, so this gene may be a lot more ancient than it appears.

We found 3 SDPs for this gene in the 2023 build of PeptideAtlas and there are five in the 2024 build. Most of the PSM are from cancer cell line experiments, but there are PSM in frontal cortex tissues and this ties in with transcriptomics support for *LIPT2-AS1* in limited brain tissues. In addition, the gene model predicted by the UniProtKB protein is probably not quite right; we find another ATG 51 codons upstream that would complete a CENP-B N-terminal DNA-binding domain and that are even conserved in bamboo shark [Figure 6A].

**Figure 6.**
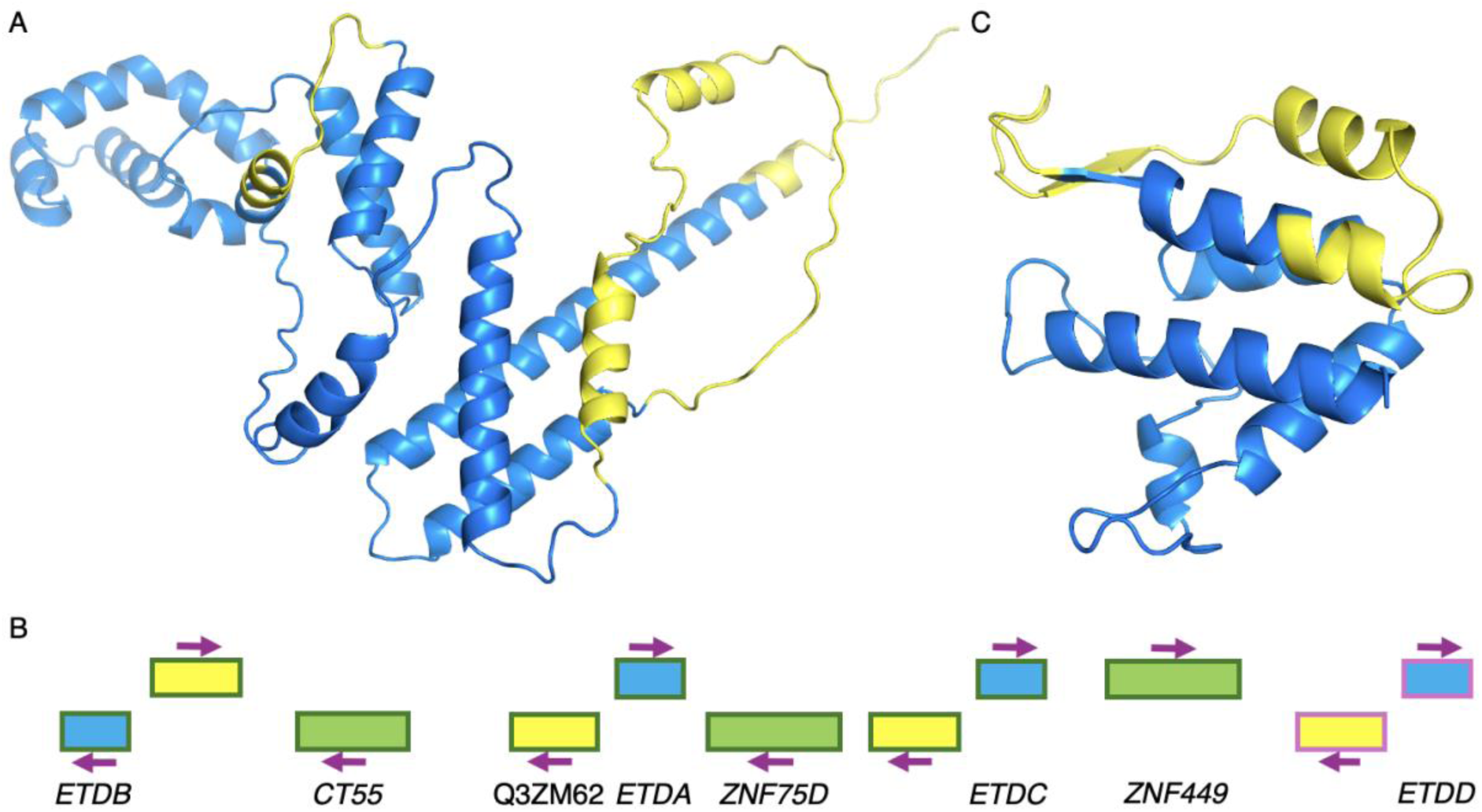
Peptides detected for *LIPT2-AS1*, *LOC107986768* and *the ETD family*. A. The AlphaFold structure of macaque *LIPT2-AS1* with the peptides detected in PeptideAtlas marked in yellow on the structure. B. A schematic representation of the human ETD family on chromosome X. ETD genes are coloured in blue, antisense ETD-like genes are shown as yellow boxes. Other intervening genes are marked in green. Gene sense is marked with arrows. Probable pseudogenes have a pink border. C. The structure of the C-terminal 134 residues of *LOC107986768* from the HHPRED server [58] with the peptides detected in PeptideAtlas marked in yellow on the structure.

UniProtKB TrEMBL protein Q3ZM62 is not annotated as an ORF in either the Ensembl/GENCODE or the RefSeq reference set. The ORF would overlap a section of 3’ UTR in *ZNF75D* just downstream of *ETDA* on chromosome X [Figure 6B]. The peptides in the most recent build of PeptideAtlas cover the protein sequence from residues 2 to 73, but the sequence of Q3ZM62 is incorrect because it would require a change in the reading frame beyond codon 80. Q3ZM62 has just over 40% identity to *ETDA*.

In fact, both *ETDA* and the Q3ZM62 ORF have three paralogues each and all on are chromosome X [Figure 6B]. As with the Q3ZM62 ORF, each of the Q3ZM62 paralogues are antisense to an EDT paralogue (*ETDB*, *ETDC*, and an unannotated ETD pseudogene). This suggests that one of the Q3ZM62 ORFs duplicated from an ETD family member, and the remaining paralogues are the result of segmental duplications of the original pair of genes. The original ETD gene duplication occurred prior to the primate clade because there is an orthologue of the Q3ZM62 ORF in mouse, though it is not antisense to mouse *Etda*. Two segmental duplications appear to have involved *ETDA* and the Q3ZM62 ORF, one pair of genes (possibly the pair involving the *ETDC* gene) duplicated right at the base of primates, the second duplication (the *ETDB* pair) is a human specific duplication. The *ETDD* pair duplicated from the *ETDC* pair after the split between old world and new world monkeys. *ETDD* is a pseudogene in great apes and its Q3ZM62-like ORF appears to be a pseudogene outside of great apes. *ETDA*, *ETDC* and Q3ZM62 are clearly protein coding across primates, but evidence for the other three ORFs is not quite so clear. In any case, all four ETD-like ORFs have been annotated as coding genes by the GENCODE annotators.

The aberrant expression patterns of *LINC00839* in cancer tissues and cell lines have been well documented [59]. The reading frame of this ORF is not conserved beyond human and all 12 peptides in PeptideAtlas are detected solely in cancer cell lines. This is a clear case of aberrant translation.

The entry HCA25a (hepatocellular carcinoma-associated antigen 25a) does not map to the human genome, but the central region that is identified by SDPs does map to multiple regions (14 in total) across the genome. This region of the HCA25a sequence is highly similar to the theoretical protein sequences of oesophageal carcinoma antigens [60]. Peptides are observed in cancer cell lines and in all likelihood whichever ORF is produces these peptides is likely to be undergoing aberrant translation. HCA25a has been tagged as unplaced because we do not know which ORF the peptides belong to.

### Possible novel ORFs not in UniProtKB

Six ORFs with peptide evidence were novel to GENCODE and not in any UniProtKB database. *LOC107986768* was supported by four SDPs, the others (*LNCPRESS1, LOC124905049, MHENCR, MNX-AS1*, and an ORF in the 3’ UTR of *PTPN18*) had two SDPs each. We believe that two of these ORFs may code for proteins in normal tissues.

*LOC107986768* is a retroviral GAG-30 derived ORF on the opposite strand from the *SCIN* coding gene and is annotated by RefSeq as coding. RefSeq notes that expression is restricted to placenta, and all the peptides in PeptideAtlas were detected in placental tissue. In the 2024 build of PeptideAtlas there are 11 peptides (5 strong and 6 non-tryptic) for this gene and all peptides map between residues 140 and 270 [Figure 6C]. This strongly suggests that there may be translation from a downstream start site, but if this is the case, it would have to be non-canonical. The reading frame is undisturbed among great apes, but if the downstream start site were used it would be undisturbed across old world monkeys. Another two novel ORFs, *PLAC4* and *ERVFRD-2*, are also retroviral and appear to be expressed mostly or solely in placenta.

*LNCPRESS1* is another retroviral ORF, this time derived from a L1 transposon. The ORF covers less than half of the original LINE-1 ORF, but what is left is largely maintained across old world monkeys. All the peptides are detected in embryonic stem cells, which makes sense given that extant LINE-1 ORFs are expressed in the early stages of development [44]. At the same time, *LNCPRESS1* RNA has been shown to be mainly nuclear and it has been postulated that the *LNCPRESS1* transcript (not protein) is a crucial part of maintaining pluripotency of cells [61]. *MHENCR* is a melanoma oncogene [62]. Peptides are detected in cancer cells and cell lines, with the majority of observations in HLA proteomics experiments. The combination of these two facts suggests that the translation of *MHENCR* is aberrant.

## Discussion

We have carried out an in-depth analysis of PeptideAtlas peptides that do not map to Ensembl/GENCODE coding genes. With the peptides that did not map to the GENCODE reference set we identified more than 250 alternative isoforms, almost 100 translated upstream regions and 34 possible coding genes that are not currently annotated by Ensembl/GENCODE. These novel coding regions are currently being analysed by GENCODE curators.

A large majority of the almost 20,000 peptides that do not map to GENCODE coding genes are most likely to be amino acid variants or post-translational modifications of peptides from known coding genes rather than novel coding regions. These peptide variants map to immunoglobulins, major histocompatibility complex proteins or other proteins known to have many variants. Many map erroneously to pseudogenes of known coding genes [33].

One surprising finding was that there were many valid PSMs for non-human proteins in the PeptideAtlas human build. The TPP identifies peptides for vertebrate, invertebrate and microbial proteins that were erroneously annotated as human by UniProtKB. The microbial proteins, particularly those of *Mycoplasma hyorhinis*, probably need to be added to the list of contaminants. We also found valid peptides for fruit fly, mouse and pig proteins in the PeptideAtlas build. While it is possible to imagine insect proteins as floating contamination similar to other contaminants such as keratin and wool, mouse and pig proteins cannot possibly be airborne. In addition, although tropomyosin is a common, ubiquitous cytoskeletal protein, there is nothing special about the mouse proteins that are misannotated as human and identified in experiments. They include a geranylgeranyl transferase, a PIDDosome adapter protein, and a substrate adapter for ufmylation. This means that the most likely explanation for the detection of these mouse proteins is large-scale cross-contamination or some kind of sample mix up.

Although we validated PeptideAtlas peptides for 34 genes that are not in the GENCODE reference set, none of them can be regarded as novel discoveries since entries in the PeptideAtlas THISP database have all been discovered previously. Just over half the entries that we found peptides for are annotated by UniProtKB as part of their human reference proteome, while another two are produced by genes annotated as coding by RefSeq. Seven of the 34 entries that we detected peptides for were identified in the large-scale ribosome profiling experiments. *MSL3P1*, *TSPY26P*, *TXLNGY*, *ZNF252P* and *LNCPRESS1* were detected by Chen *et al* [18], while *LIPT2-AS1, and AARSD-AS1* were reported by Van Heesch *et al* [19].

We analysed the PSMs, tissue expression patterns and cross-species conservation for all 34 genes in this analysis. We believe that almost half are likely to be coding genes, including 8 of the 18 genes from the UniProtKB human proteome and 5 of the 8 proteins from the UniProtKB TrEMBL database. Most of these 16 likely coding genes appear to be recent evolutionary innovations, but still have clear evidence of coding conservation. This coding conservation is clear even in those ORFs that have novel stop codons in great apes (*ERVFRD-2*, *LIPT2-AS1*).

We believe that 14 of the ORFs are probably producing peptides as a result of aberrant translation. These ORFs have peptides detected only in HLA proteomics experiments (*LINC03040*), only in cancer cell experiments (*ZNF252P*), or are known to be cancer antigens (*MHENCR*). The human reading frames of 12 of these 14 ORFs are not conserved in any species and the other two are not conserved beyond great apes. Six are already described as cancer relevant genes.

The remaining 4 proteins with multiple SDPs were unplaced in the genome. *FAM230A*, Q9UF83, HCA25a and *POM121L1P* mapped to multiple regions in the human genome, and it would be almost impossible to decide which of the regions was being translated. Two of these entries only had peptides in cancer cell line experiments, but almost all peptides for *FAM230A* are found in testis or sperm, and there are experimental peptides from sperm for *POM121L1P*, though the PSMs are poor for these peptides. Although something similar to the *POM121L1P* peptides may be sperm expressed, there are more than 150 possible different POM121 pseudogene locations in the human genome, scattered mostly across chromosomes 5, 6 and 22.

LINE-1 ORF1 could conceivably be added to the list of unplaced ORFs. As with the POM121 pseudogenes, LINE-1 ORF1 is present in multiple copies in the genome. It is also the unannotated ORF with most peptide evidence in PeptideAtlas, and almost all peptide-spectrum matches are from cancer cells or cell lines [21]. Between LINE-1 ORF1 and the unplaced and likely aberrant coding genes there were 17 unannotated non-conserved ORFs that are supported only by peptide evidence from cancer cells or cell lines, showing that aberrant translation products may be common in cancer cells. This adds to the evidence of the substantial dysregulation of translation in cancer cells [63–65].

The peptides for novel ORFs that have no cross-species support and expression limited to cancer cells illustrates what may become a whole new level of complexity in the annotation of coding genes if many of these ORFs are present in ribosome profiling experiments. The first question is whether the proteins produced by these ORFs have functional roles. Despite the peptides, the conservation evidence suggests that this is unlikely, but it cannot be ruled out completely. The second question is, if these proteins are not functionally important, how should they be annotated? Should they be added to the reference gene set as coding genes for the sake of completeness, as is currently the case with genes like *HMHB1* [66] and *MYEOV* [67], or should they be labelled separately so that they cannot be mistaken for genes that produce biologically relevant proteins?

